# *Gpnmb*-positive lipid-associated macrophages contribute to lipid clearance during luteolysis

**DOI:** 10.64898/2026.08.31.748418

**Authors:** Yu-Hsiang Liao, Kashish Jain, Yi A Ren

## Abstract

Macrophages are important regulators of ovarian physiology, yet the complexity in their subpopulations and functions is far from fully understood and is emerging from single-cell analysis of diverse physiological and pathological contexts. One open question in ovarian physiology is how massive lipid loads during structural luteolysis are resolved without inducing inflammation. Here, by integrating single-cell and spatial transcriptomic datasets from mouse and human ovaries with high-resolution immunostaining and whole-mount imaging, we identified a *Gpnmb*+ subset of macrophages that dynamically expand during luteolysis in the mouse ovary, and a similar *GPNMB*+ macrophage population is also present in human ovarian scRNA-seq datasets. Transcriptomic profiling revealed that these *Gpnmb*+ macrophages resemble lipid-associated macrophages (LAMs) in adipose tissues in obesity, displaying strong enrichment in phagocytosis, lysosomal lipid processing, and reverse cholesterol transport. Notably, unlike phagocytic macrophages that only clear cellular debris, *Gpnmb*+ macrophages represent a distinct, specialized subset dedicated to processing lipid-rich targets: they selectively infiltrate regressing corpora lutea (CL), internalizing lipid droplets and interacting with dying luteal cells via apoptosis (*Thbs1-Sdc1/4*), efferocytosis (*Pros1*-*Mertk*), and cell recruitment (*Cxcl12/Cxcr4*) signaling axes. *Gpnmb*+ macrophages also surround oocytes in atretic follicles. Through integrated transcriptomic and imaging analyses, our data suggest that *Gpnmb*+ macrophages execute a complete cascade of multi-stepped lipid processing, including lipid droplet internalization, lysosomal and lipophagic degradation, and ABCA1/ABCG1-mediated cholesterol efflux. Furthermore, whole-mount imaging revealed close physical interaction between *Gpnmb*+ macrophages and LYVE1+ lymphatic endothelial cells, supporting a non-inflammatory resolution for lipid clearance via lymphatic circulation. Collectively, our findings establish *Gpnmb*+ macrophages as specialized LAMs essential for maintaining cyclic ovarian lipid homeostasis. This work redefines the paradigm of LAM biology by demonstrating their vital homeostatic role in normal ovarian physiology, bridging immune-mediated tissue remodeling and reproductive physiology.

## Introduction

Macrophages play a wide range of roles in ovarian physiology, such as primordial follicle activation, ovulation, and vascular integrity in corpus luteum (CL) [1–5]. Macrophages are abundant in adult mammalian ovaries. They are located in the ovarian stroma throughout the estrus cycle in mice, with their density increasing from antral to ovulating follicles [6]. While they are not present in granulosa cell (GC) layers in healthy follicles before ovulation, macrophages are found in both theca and granulosa cell layers of atretic follicles and execute phagocytosis [6, 7]. After ovulation, macrophages are found in the forming CL, and contribute to angiogenesis in CL [8]. As the CL gradually degenerate (luteolysis), more macrophages infiltrate them [6] for the clearance of luteal cells.

Luteolysis is triggered either by a reduction in luteotropic support or by an elevation of the luteolytic signal, prostaglandin F2α (PGF2α). While the former mechanism predominates in humans and non-human primates, the latter serves as the primary driver in mice and livestock species [9, 10]. Functional luteolysis is characterized by a reduction in progesterone production, occurring approximately 2 days after the luteinizing hormone (LH) surge in mice, whereas structural luteolysis is marked by luteal cell death and tissue clearance, starting from around 3 to 4 days post-LH surge in mice [11]. Immune cells are well established as key regulators in initiating luteal regression and executing subsequent tissue clearance [6]. CL represent one of the most active steroidogenic tissues in the body and accumulate abundant lipid droplets rich in triglycerides and cholesterol [12–14]. However, the precise mechanisms by which cells clear these excess lipids during luteolysis remain poorly understood. While luteal cells may efflux cholesterol via ATP-binding cassette transporters A1 (ABCA1) during luteolysis [15], whether immune cells are involved, and how the local ovarian microenvironment processes and clears this excessive amount of lipids without inducing chronic lipotoxicity or inflammation remains a fundamental, unresolved question in ovarian biology.

Historically, functional insights into ovarian macrophages have primarily relied on observation, macrophage depletion strategies, and binary classifications, such as the classical proinflammatory (M1) and alternatively activated (M2) macrophages [16]. While M1 and M2 macrophages have been implicated in distinct processes like folliculogenesis [17] and primordial follicle activation [1], these classifications may not capture the microenvironmental adaptations of specialized tissue-specific macrophages. Recent advances in single-cell RNA sequencing (scRNA-seq) have enabled the discovery of specialized macrophage subpopulations in various biological contexts. For instance, Triggering Receptor Expressed on Myeloid Cells 2 (*Trem2*)-and Glycoprotein nonmetastatic melanoma protein B (*Gpnmb*)-expressing lipid-associated macrophages (LAMs) have been identified in obese adipose tissue, fatty liver, and neurodegenerative disorders, where they act as essential homeostatic sensors that internalize and metabolize excessive lipid [18]. Furthermore, a recent study has highlighted *Gpnmb* as a marker of multinucleated giant cells (MNGCs) and fibro-inflammatory remodeling in the aging ovary [19], suggesting that *Gpnmb*+ macrophages accumulate in response to age-associated tissue remodeling. Although a recent study reported the presence of *Gpnmb*+ macrophages in normal cycling mouse ovaries [20], their physiological role, spatiotemporal dynamics, and lipid-processing capabilities remain unexplored.

In this study, to determine whether *Gpnmb*+ macrophages function as a specialized, LAM-like population that plays a role in lipid metabolism during normal ovarian homeostasis, we integrate public mouse [21, 22] and human ovarian scRNA-seq datasets [23] with spatial transcriptomics [24], and spatiotemporal immunostaining. By integrating these data, we aim to characterize their functional contributions to lipid processing and tissue remodeling in normal ovarian physiology.

## Results

### Identification of an estrus-enriched *Gpnmb*+ macrophage population in the mouse ovary

To investigate the function of macrophages in ovarian physiology, we initially analyzed published scRNA-seq datasets of the cycling mouse ovary [21]. Macrophages for downstream analysis were selected based on annotation in the original paper (markers include *Csf1r* and *Ly86*, Fig. S1A) [21]. Subsequent unsupervised clustering resolved these macrophages into seven distinct subclusters (Fig. 1A), including one with enriched transcripts of *Gpnmb*. The marker genes of each subset are provided in Supplementary File 1. Analysis of subclusters of macrophages across the four estrous stages (proestrus, estrus, metestrus, and diestrus) demonstrated stage-specific enrichment of individual subclusters (Fig. 1B, left). Notably, the *Gpnmb*+ subcluster exhibited an expansion in the estrus phase (Fig. 1B, left). Uniform Manifold Approximation and Projection (UMAP) plot also showed an enrichment of cells in the estrus stage within *Gpnmb*+ macrophages (Fig. 1A, right, and Fig.1B, right). Analysis of this dataset also demonstrated that transcripts of *Gpnmb* were predominantly in macrophages in the mouse ovary (Fig. S1B). To validate these findings, we analyzed an independent ovarian scRNA-seq dataset from young and aged mice [22]. Consistently, re-clustering of the “tissue macrophage” population (annotated using *C3ar1*, *Itga9*, and *C5ar1* as markers) from both ages recapitulated the presence of a *Gpnmb*+ macrophage subcluster (Fig. S2A & B), aligning with a recent report of *Gpnmb*+ macrophages in normal cycling ovaries [20].

**Figure 1.**
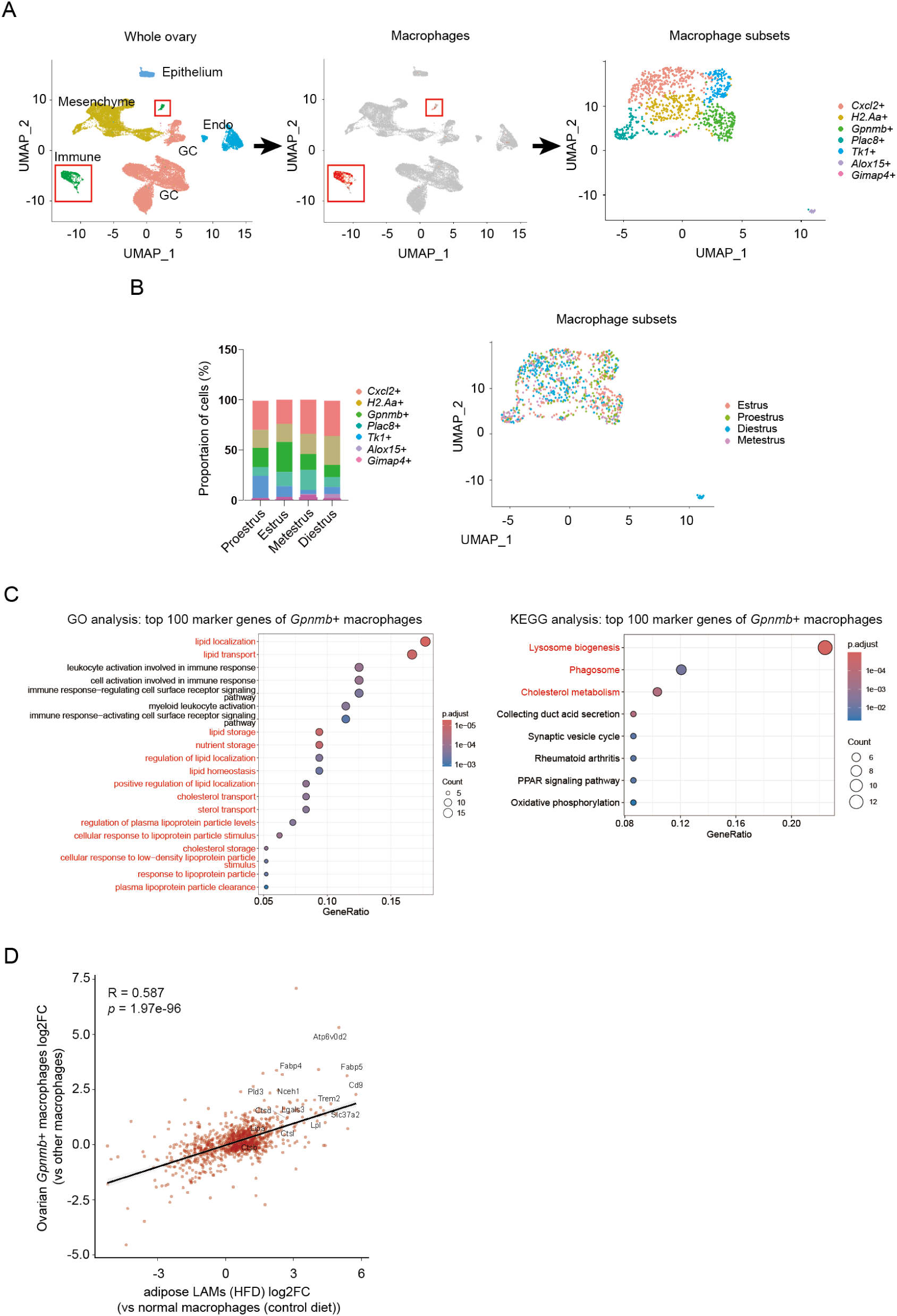
Identification of *Gpnmb*+ macrophages in mouse ovarian scRNA-seq dataset. (A) Uniform Manifold Approximation and Projection (UMAP) of major cell types (left), macrophages (middle), and subsets of macrophages (right) in the ovary. (B) Cell proportion of macrophage subsets at different stages of the estrous cycle (left). UMAP showing the distribution of macrophages at different stages of the estrus cycle (right). (C) Gene ontology (GO) analysis and Kyoto Encyclopedia of Genes and Genomes (KEGG) analysis of top 100 marker genes of *Gpnmb*+ macrophages. (D) Transcriptomic comparison between ovarian *Gpnmb+* macrophages and LAMs in adipose tissue. Scatter plot showing differentially expressed genes (DEG, log_2FC) of ovarian *Gpnmb+* macrophages (y-axis, compared with other ovarian macrophages) and LAMs in adipose tissue (x-axis, compared with normal macrophages in normal adipose tissue) [18]. Data and analyses in Figure 1 were based on an ovarian scRNA-seq dataset [21].

To investigate the function of *Gpnmb*+ macrophages, we performed Gene Ontology (GO) and Kyoto Encyclopedia of Genes and Genomes (KEGG) pathway analyses based on the differentially expressed genes (DEGs) between *Gpnmb*+ macrophages and other subsets of macrophages in the ovary. *Gpnmb*+ macrophages displayed robust enrichment in pathways related to phagocytosis and lipid metabolism (Fig. 1C), which are also major functions of lipid-associated macrophages (LAMs) found in obese adipose tissue [18]. The resemblance between ovarian *Gpnmb*+ macrophages and adipose LAMs is further demonstrated by a shared transcriptomic shift: namely, the difference between *Gpnmb*+ subcluster vs. other ovarian macrophages, and the difference between adipose LAMs from obese mice vs. adipose macrophages from normal-weight mice. We observed a robust positive correlation (computed based on the trend and fold-change of differences of differentially expressed transcripts) between the two differential gene profiles (Fig. 1D), Collectively, these data support the identification of a localized, LAM-like macrophage population in the mouse ovary.

### Cross-species conservation of the *GPNMB*+ ovarian macrophage population between mouse and human

To determine whether the *Gpnmb*+ macrophage population and their function in murine ovaries are evolutionarily conserved in humans, we compared the scRNA-seq data from mouse ovaries with previously published scRNA-seq datasets from human ovaries derived from three healthy donors (aged 28, 37, and 31) [23]. Unsupervised clustering of human ovarian macrophages (identified by the co-expression of *CD68* with *CD14* or *FOLR2* [23], Fig. S3A) resolved four distinct macrophage subpopulations, including a *GPNMB+* cluster (Fig. 2A). The marker genes of each subset are provided in Supplementary File 2. Next, to assess the evolutionary conservation of this *GPNMB+* cluster across species, we cross-referenced the respective marker genes of *Gpnmb+/GPNMB+* macrophages from mice and humans. Among the 177 marker genes in the mouse *Gpnmb*+ cluster and 83 marker genes in the human *GPNMB*+ cluster, we identified an overlap of 22 genes, including *GPNMB/Gpnmb*, *TREM2/Trem2*, *FABP5/Fabp5*, and *LIPA/Lipa* (Fig. 2B), which were selectively enriched in *GPNMB+*/*Gpnmb+* macrophages across human and mouse UMAPs (Fig. 2C). The significance of this overlap was evaluated via a hypergeometric test using the R package Phyper, which confirmed a highly significant correlation between the mouse and human clusters (*p* =1.931795e-19) (Fig. 2B). Subsequent Gene Ontology (GO) analysis on these 22 conserved genes revealed enrichment in functional pathways critical for metabolic homeostasis, including lipid transport, tissue remodeling, and cholesterol efflux (Fig. 2D). The pathways related to “tumor necrosis factor production” suggest the anti-inflammatory function of the *Gpnmb*+ macrophages, as the genes listed in the pathways (*Igf1/Trem2/Acp5/Gpnmb/Cd84*) have been shown to inhibit inflammation [25, 26]. Taken together, these comparative transcriptomic analyses demonstrate that the molecular signature and function of ovarian *Gpnmb*+ macrophages are highly conserved between mouse and human.

**Figure 2.**
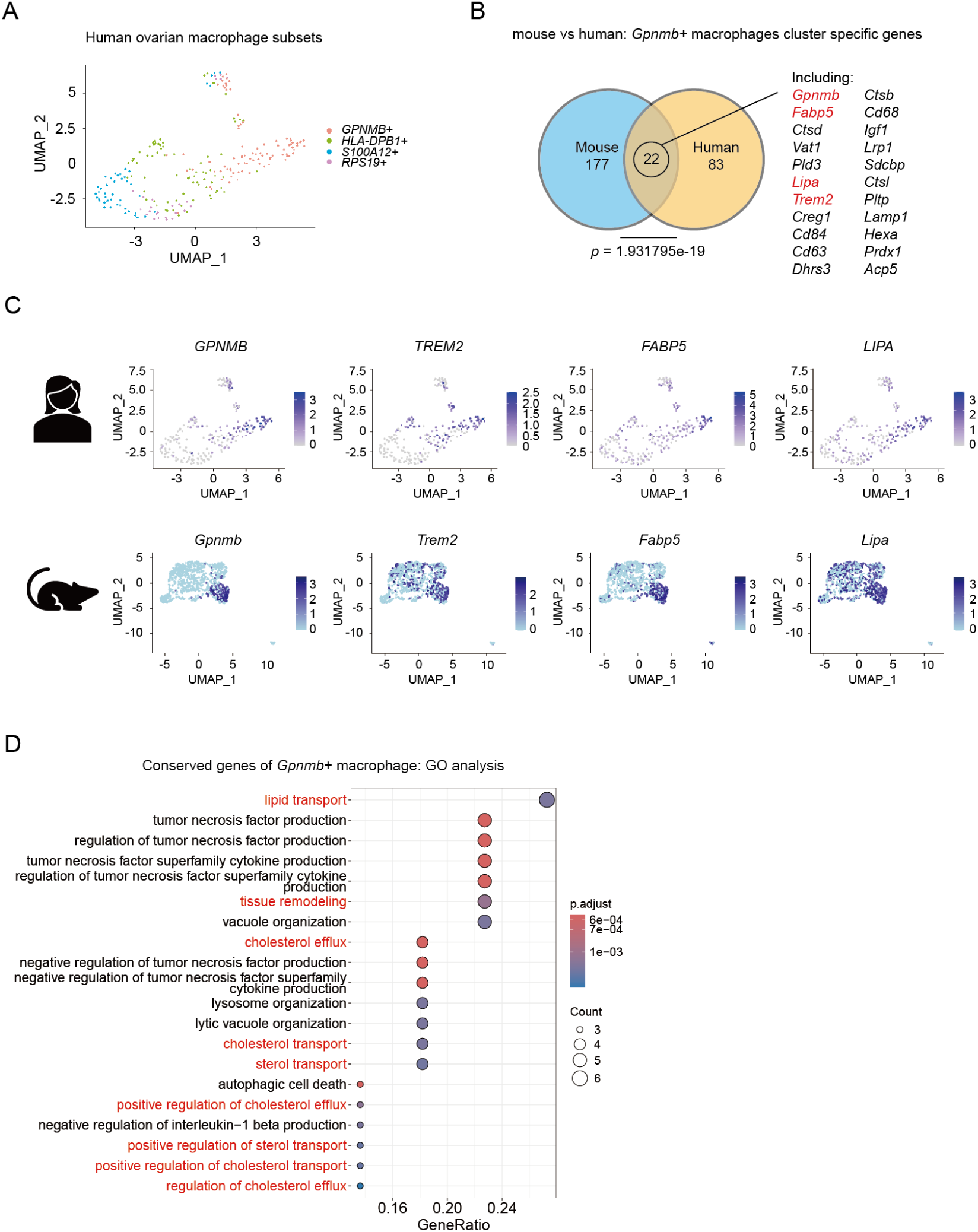
*Gpnmb*+ subset of macrophages in the ovary is conserved between human and mice. (A) UMAP of macrophage subsets in a scRNA-seq dataset from human ovaries [23]. (B) A Venn diagram illustrating the overlap of significantly expressed genes (*p* < 0.05) in *Gpnmb*+ macrophages between mouse [21] and human ovaries [23]. To facilitate the analysis, human gene orthologs were converted to their corresponding mouse gene nomenclature. Statistical significance of the overlap was assessed via a hypergeometric test using the phyper function in R, revealing a highly significant enrichment between the mouse and human datasets (*p* = 1.931795e-19). (C) UMAPs showing the expression of *GPNMB/Gpnmb, TREM2/Trem2, FABP5/Fabp5,* and *LIPA/Lipa* within human (top) and mouse (bottom) ovarian macrophage clusters. (D) GO analysis of the 22 conserved genes between *Gpnmb+ macrophages* from mouse and human ovary.

### Spatiotemporal analysis identifies *Gpnmb*+ macrophages in atretic follicles and regressing corpora lutea

To gain a comprehensive understanding of where *Gpnmb*+ macrophages are located in the ovary, we first analyzed spatial transcriptomic data of ovaries from superovulated immature mice over multiple time points during the preovulatory stage before the formation of CL (Fig. 3A) [24]. We found that the number of *Gpnmb*+ cells was higher in the immature stage compared to the preovulatory stage, except for 4h post-hCG, when *Gpnmb*+ cells formed a large cluster in the ovarian stroma (Fig. 3B & 3C). Across all time points examined, *Gpnmb*+ cells appeared to be enriched in atretic follicles, preantral follicles, or to form clusters in the stromal region (Fig. 3C). As there were significantly more atretic follicles in ovaries from immature mice compared to other time points, the enrichment of *Gpnmb*+ cells in these structures explains their higher number in the ovaries from immature mice.

**Figure 3.**
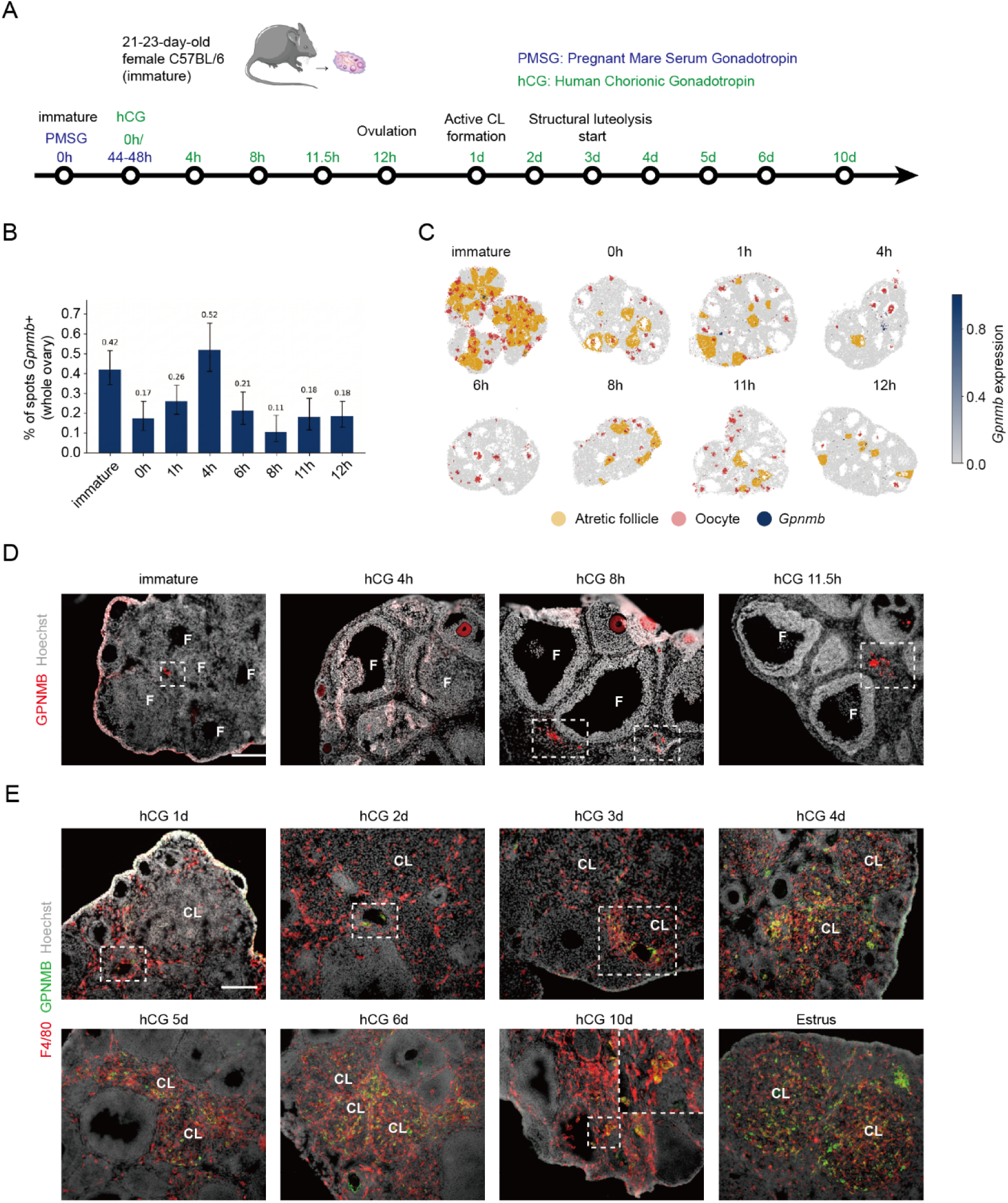
Ovarian *Gpnmb*+ macrophages surround the oocytes in atretic follicles and are located in regressing corpora lutea. (A) A schematic of the timeline of sample collection relative to superovulation in immature mice. (B) The percentage of *Gpnmb*+ cells in whole ovary section at different time points after hCG injection. (C) The distribution of *Gpnmb*+ cells, atretic follicles, and oocytes in the spatial transcriptomic dataset. (D) Immunostaining of GPNMB in ovaries collected from immature mice either without superovulation stimulation or at 4, 8, and 11.5 hours following hCG administration. Square: *Gpnmb*+ macrophages. Scale bar: 200um. (E) Representative images of immunostaining of GPNMB and F4/80 in ovaries collected from mice at 1, 2, 3, 4, 5, 6, and 10 days post-hCG and in ovaries of natural cycling mice (estrus). Square: *Gpnmb*+ macrophages. Scale bar: 200um. Data and analyses in B and C were based on an ovarian spatial transcriptomic dataset [24].

To confirm the localization of *Gpnmb*+ macrophages in the mouse ovary as revealed by spatial transcriptomics (Fig 3C), we performed immunostaining of GPNMB on ovaries from superovulated immature mice at multiple preovulatory time points (Fig. 3A). The result revealed that *Gpnmb*+ cells were located within smaller follicles that appear to be degenerating, and their number was relatively low across the ovary. This low abundance is consistent with the general macrophage population dynamics following PMSG-mediated follicular rescue from atresia, prior to the first ovulation and CL formation (Fig. 3D) [27]. Notably, although some GPNMB signals were detected in oocytes, we considered it non-specific, as *Gpnmb* is predominantly expressed in macrophages (Fig. S1B) and oocytes are known to exhibit non-specific staining [28]. Based on a recent study reporting the presence of *Gpnmb*+ macrophages in the CL [20], we also tracked the spatiotemporal dynamics of *Gpnmb*+ macrophages at multiple time points post-hCG across later stages of the superovulation model. At 1 day (1d) post-hCG (active CL formation), macrophages (labeled with F4/80) infiltrated the newly formed CL but remained immunonegative for GPNMB inside CL. During this stage and up to 2d post-hCG, *Gpnmb*+ macrophages were still restricted within smaller follicles (Fig. 3E). A distinct shift occurred at 3d post-hCG, the onset of luteolysis in normal cycling mice [11], where we observed the first appearance of *Gpnmb*+ macrophages inside CL (Fig. 3E). By 4d to 6d post-hCG, as structural luteolysis progressed, a substantial influx of *Gpnmb*+ macrophages occurred, indicated by their high abundance within the regressing CL. At 10d post-hCG, we observed fewer *Gpnmb*+ macrophages in the CL, and some of them were confined to structures that appeared to be degenerated CL (Fig. 3E). Because structural luteolysis takes place around 3-4 days after the LH surge [11], and this coincides with the subsequent LH surge and ovulation, the high abundance of *Gpnmb*+ macrophages around 3-4 days post-hCG matches the estrus-specific enrichment of *Gpnmb*+ macrophages observed in scRNA-seq datasets (Fig. 1B). Taken together, the precise alignment between *Gpnmb*+ macrophage infiltration and structural luteolysis (peaking from 4d to 6d post-hCG/LH surge) supports their primary function in clearing regressing luteal tissue.

### *Gpnmb*+ macrophages in atretic follicles are a specialized subset of phagocytic macrophages

These temporal dynamics were further validated by qPCR of ovaries from immature mice before and after superovulation: while transcript levels of ovarian *Gpnmb* and *Trem2* remained low from immature stages to 2d post-hCG (48h post-hCG), their transcript levels exhibited a multifold upregulation at 5d post-hCG (Fig. 4A). Interestingly, the transcript level of *Atp6v0d2*, a key lysosomal gene [29], was high at both immature stage and 5d post-hCG (Fig. 4A). Given that *Atp6v0d2* serves as a marker for active lysosomal phagocytosis and is expressed only by macrophages in the ovary (Fig. 4B), its high transcript level indicates that the subset of macrophages that have phagocytic machinery to clear apoptotic cellular debris is present both in atretic follicles (which predominate in immature ovaries) and regressing CL (which predominate in hCG 5d). In contrast, the notable upregulation of *Gpnmb and Trem2* only at 5d post-hCG suggests that *Gpnmb*+ macrophages do not only participate in generalized lysosomal phagocytosis of cellular debris (as *Gpnmb*+ macrophages also express *Atp6v0d2*, Supplementary File 1), but are also functionally specialized to manage lipid-rich microenvironments. Intriguingly, spatial transcriptomic data indicate that *Gpnmb*+ cells do not represent all macrophages in the atretic follicles (Fig. 4C). Indeed, among all time points analyzed, *Gpnmb*+ macrophages within atretic follicles consistently and selectively surrounded oocytes (Fig. 4D & S4A), which contained more intracellular lipids compared to surrounding GCs [30], supporting the specialized role of *Gpnmb*+ macrophages in processing lipid-rich cellular targets (Fig. 4D & S4A). Notably, while immunostaining of the pan-macrophage marker (F4/80+) showed a broad presence of total macrophages within atretic follicles (Fig. 4E, white arrow), the *Gpnmb*+ subset exclusively appeared around the oocytes (Fig. 4E, asterisk). Taken together, these data suggest that *Gpnmb*+ macrophages are a specialized phagocytic subset with lipid-processing capacity.

**Figure 4.**
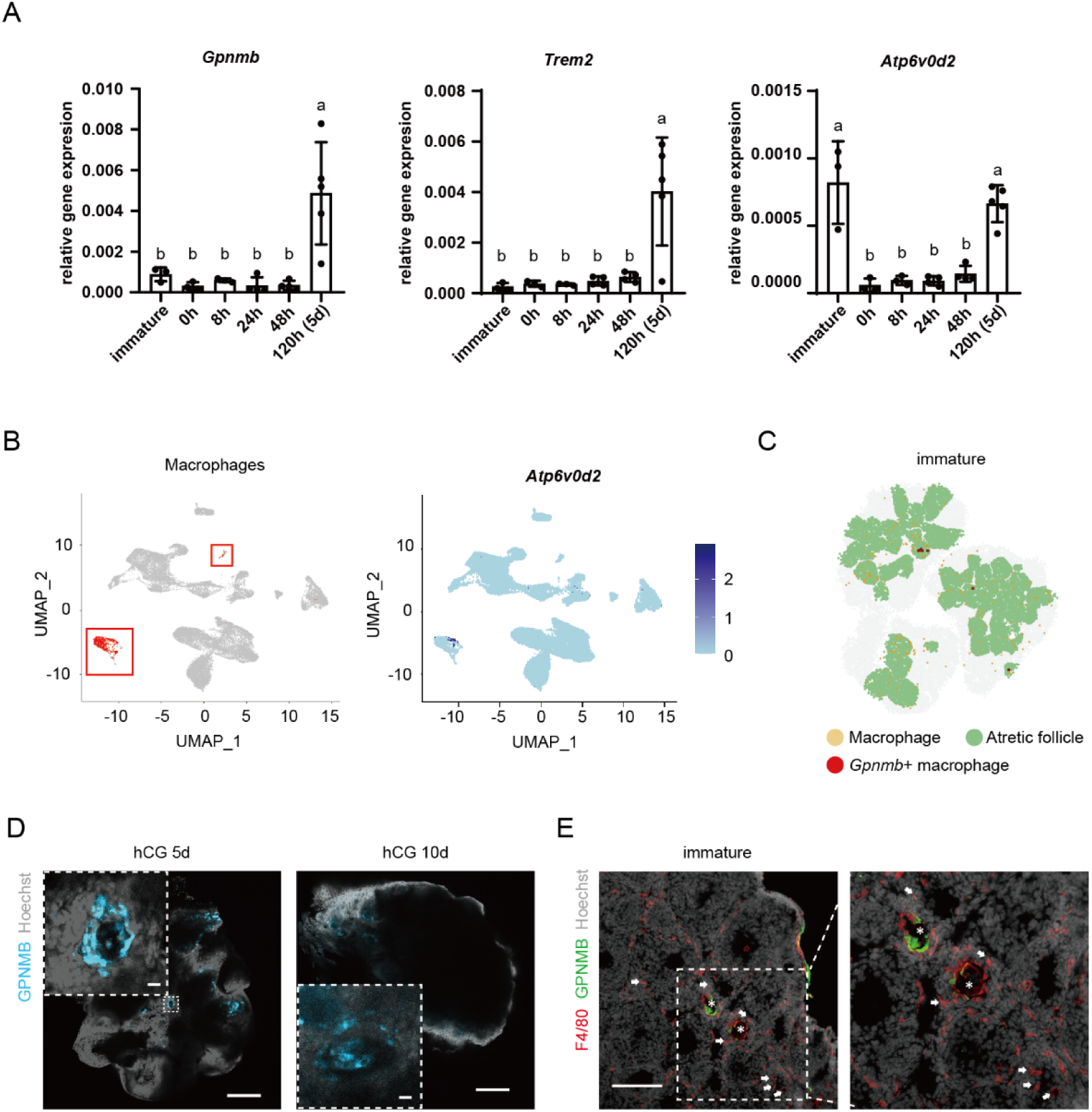
Ovarian *Gpnmb*+ macrophages are a specific subset of phagocytic macrophages in atretic follicles. (A) RT-qPCR analysis of *Gpnmb, Trem2, and Atp6v0d2* mRNA levels in mouse ovaries before and after superovulation (n = 3-5). *Rpl19* served as an internal control. Different letters represent significant differences determined by one-way ANOVA followed by Tukey’s multiple comparison. *p* < 0.05. (B) UMAP showing that the expression of *Atp6v0d2* is restricted in macrophages in the mouse ovary. (C) The distribution of *Gpnmb*+ macrophages in the spatial transcriptomic dataset showing that they did not represent all the macrophages in the atretic follicles. (D) One section of whole-mount staining of GPNMB in ovaries collected from mice at d5 and 10 following hCG administration showing that *Gpnmb*+ macrophages surround the oocytes. Scale bar: 200um (big). Scale bar: 10um (small). (E) Immunostaining of GPNMB and F4/80 in ovaries collected from immature mice showing that *Gpnmb*+ macrophages are a specific subset of phagocytic macrophages in atretic follicles. Scale bar: 200um. Data and analyses in B and C were based on an ovarian scRAN-seq dataset [21] and spatial transcriptomic dataset [24] respectively.

### *Gpnmb+* macrophages selectively interact with dying luteal cells to coordinate efferocytosis during luteolysis

To determine whether *Gpnmb*+ macrophages preferentially target regressing CL, we performed ligand-receptor cell-cell interaction analysis of scRNA-seq data between *Gpnmb+* macrophages and active (defined by *Top2a* and transcripts of steroidogenic enzymes [21]) versus dying luteal cells (defined by *Cdkn1a, Sdc4, Cldnd1,* and *Btg1* [21]) during the estrus stage. The focus on the estrus stage is based on the localization of *Gpnmb+* macrophages within the CL by immunostaining at 3-6d post-hCG (Fig. 3D) and the initial discovery of this subset enriched in estrus (Fig. 1B). Our analysis revealed a greater number of active signaling pathways between *Gpnmb+* macrophages and dying luteal cells compared to active luteal cells (Fig. 5A), supporting selective interaction with dying luteal cells. Detailed pathway mapping identified a unique signaling interaction specifically enriched between *Gpnmb+* macrophages and dying luteal cells, including pathways modulating lipid/phagocytic-sensing programs (*App*-*Cd74*, *App*-*Trem2/Tyrobp*), efferocytosis (clearing of dying cells by phagocytosis) and cellular clearance (*Pros1*-*Mertk/Axl*), cell recruitment (*Cxcl12-Cxcr4,* Semaphorin pathways), and broader structural matrix remodeling (collagen and laminin interactions) (Fig. 5B, left); conversely, feedback signaling from *Gpnmb*+ macrophages switched from pro-survival signal (*Igf1*-*Igf1r*) toward active luteal cells to degeneration-promoting signals toward dying luteal cells, including matrix adhesion (*Spp1-Itga9/Itgb1*), apoptosis/clearance induction (*Thbs1*-*Sdc1/4/Cd47*, *Tnfsf12-Tnfrsf12a*), and baseline structural interactions (*Col1a2-Sdc1/4/Itga9+Itgb1, Entpd1-Adora1*) (Fig. 5B, right). Taken together, these data support a model that *Gpnmb+* macrophages are actively recruited by and selectively interact with dying luteal cells to execute phagocytosis, thereby contributing to the regression of the CL.

**Figure 5.**
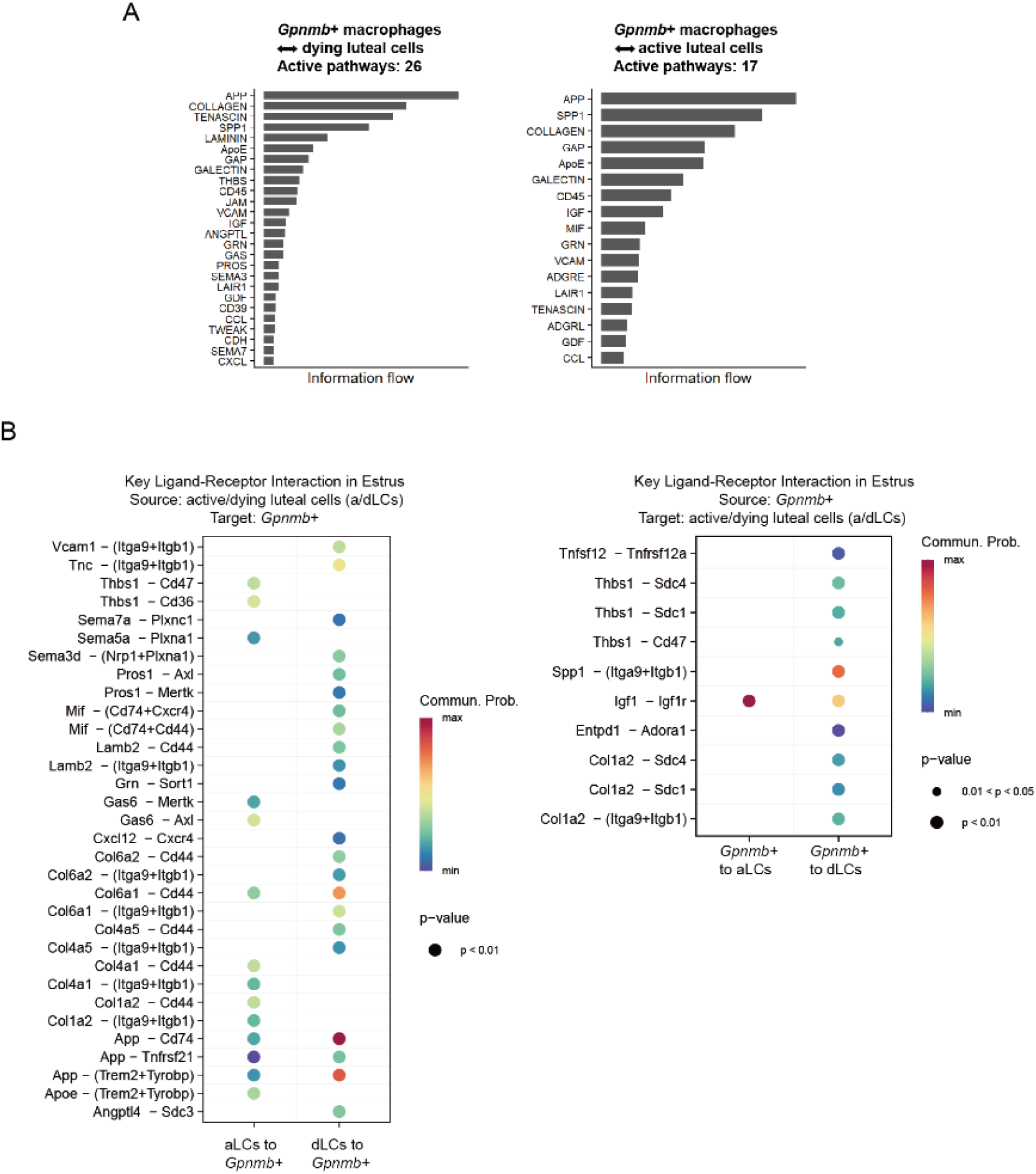
Ovarian *Gpnmb*+ macrophages interact more with dying luteal cells compared to active luteal cells. (A) Bar plots showing the major active signaling pathways between *Gpnmb*+ macrophages and either dying luteal cells (left, 26 active pathways) or active luteal cells (right, 17 active pathways) in estrus inferred by CellChat analysis [21, 61]. The pathways are ranked based on their total communication probability within each cell pair interaction. (B) Detailed key ligand-receptor interactions between *Gpnmb*+ macrophages and either dying luteal cells or active luteal cells in estrus, inferred by CellChat analysis. Left: luteal cells to *Gpnmb*+ macrophages; right: *Gpnmb*+ macrophages to luteal cells [21, 61].

### Ovarian *Gpnmb+* macrophages exhibit robust lipid uptake and lysosomal processing capabilities in vivo

In addition to the phagocytic profile, *Gpnmb*+ macrophages were enriched for transcripts associated with lipid transport and cholesterol metabolism, as shown in Fig. 6A. Therefore, we hypothesized that this population is functionally specialized to process the massive amount of lipid accumulated in luteal cells during luteolysis [12–14]. High transcript levels of *Cd36* and *Mertk* support their ability of phagocytosis and lipid uptake (Fig. 6A). To functionally assess the lipid-uptaking capacity of *Gpnmb*+ macrophages in vivo, we performed neutral-lipid staining via BODIPY across the superovulation time course. At 3d post-hCG, *Gpnmb*+ macrophages were found to be restricted to lipid-rich regions of the regressing CL (Fig. 6B). Notably, a subset of these macrophages exhibited intracellular lipid droplets starting from 4d post-hCG, suggesting the engulfment of lipid-rich debris from luteal cells. This association with lipid persisted through 6d post-hCG, and in ovaries at the estrus stage in random cycling adult mice (Fig. 6B). We further performed whole-mount immunostaining and imaging with high resolution, demonstrating the internalization of luteal lipid droplets by *Gpnmb*+ macrophages in the CL at 5d post-hCG (Fig. 6C). Together, these findings establish the capacity of *Gpnmb*+ macrophages to take up lipids in regressing CL.

**Figure 6.**
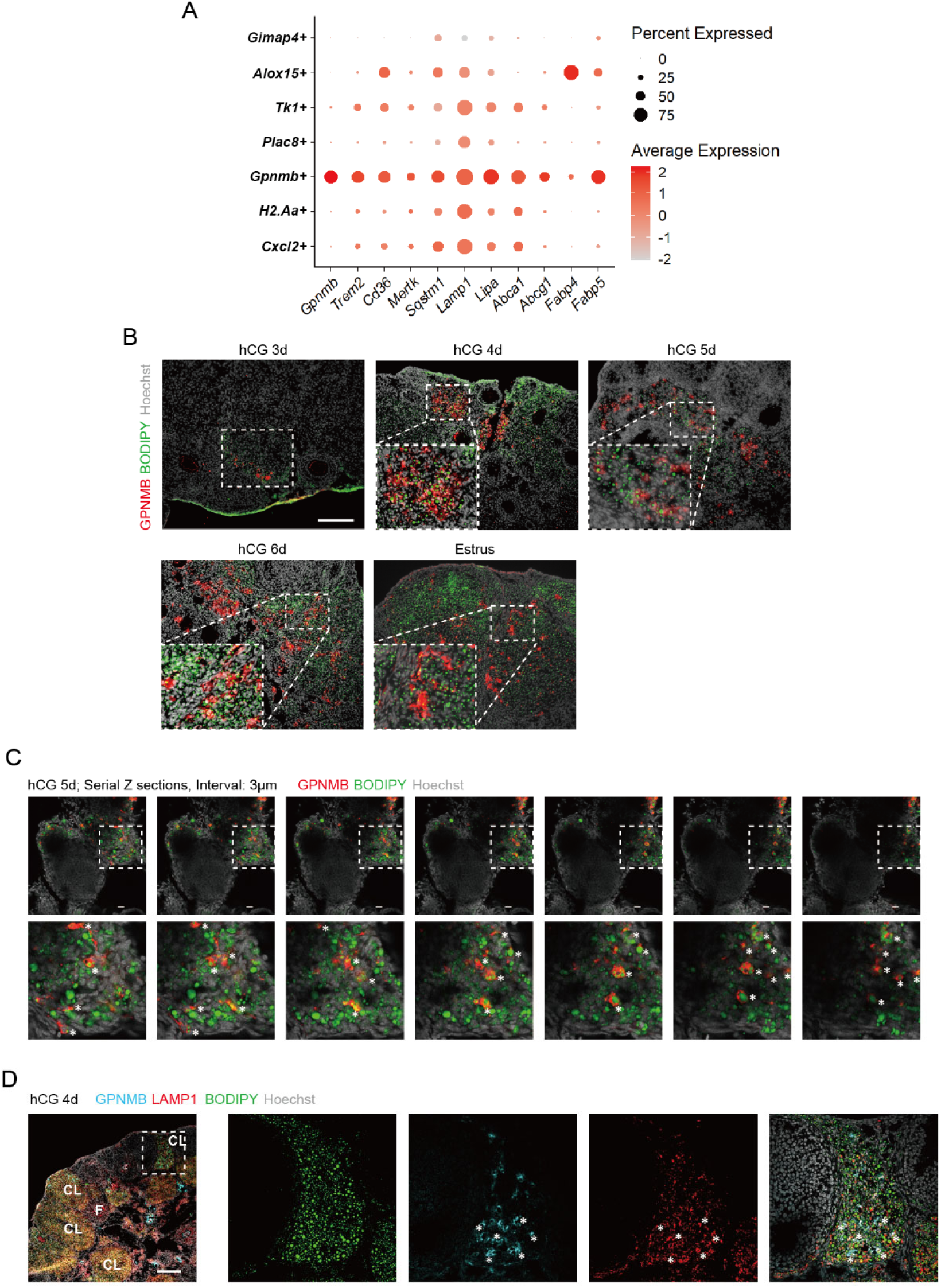
Ovarian *Gpnmb+* macrophages exhibit robust lipid uptake and lysosomal processing capabilities. (A) Dot plot of selected DEGs in ovarian *Gpnmb*+ macrophages compared with other subsets of ovarian macrophages. Gene symbols along the Y-axis represent the signatures for the seven subsets of macrophages, respectively, that we identified by analyzing an ovarian scRNA-seq dataset [21]. (B) Immunostaining of GPNMB and BODIPY in ovaries collected from immature mice at d3-6 post-hCG and in adult natural cycling mice (estrus). Scale bar: 200um. (C) Serial Z-sections of whole-mount staining of GPNMB and BODIPY in ovaries from mice at d5 post-hCG at macro (top) and micro (bottom) scales. Asterisk: *Gpnmb*+ macrophages with lipid droplets (BODIPY). Scale bar: 20um. (D) Immunostaining of GPNMB, LAMP1, and BODIPY in ovaries collected from mice at d4 post-hCG. Asterisk: *Gpnmb*+ macrophages with lipid droplets (BODIPY) and LAMP1. Scale bar: 200um.

Macrophage-mediated lipid uptake can be coupled with lipophagy, a form of autophagy selectively breaking down intracellular lipid droplets, to facilitate the degradation of lipid droplets [31]. Intriguingly, the core autophagy adaptor *Sqstm1* (encoding p62) [32] was highly enriched in *Gpnmb*+ macrophages (Fig. 6A). Because the punctate accumulation of LC3 serves as a standard indicator of autophagosome membrane assembly [33], we performed LC3 immunostaining to evaluate the presence of autophagic structures in *Gpnmb*+ macrophages. While robust LC3 immunoreactivity was detected in granulosa cells of follicles before ovulation, scattered LC3 staining was detected in CL with some colocalizing with GPNMB (Fig. S5A), indicating that lipophagy may be a mechanism for lipid processing in ovarian *Gpnmb+* macrophages. Additionally, high transcript levels of *Lamp1* and *Lipa* indicated active lysosomal degradation of lipid (Fig. 6A). To assess this idea, we co-stained BODIPY, LAMP1 (the marker of lysosome), and GPNMB. Indeed, we observed BODIPY staining colocalized with LAMP1 in *Gpnmb+* macrophages through 4d to 6d post-hCG, indicating active lysosomal processing of the engulfed lipids (Fig. 6D). Finally, the high transcript levels of *Abca1*, *Abcg1, Fabp4,* and *Fabp5* support the cholesterol efflux [34] and fatty acid transportation ability of *Gpnmb+* macrophages (Fig. 6A). Collectively, integrated transcriptomic profiling and immunostaining characterize the capacity of *Gpnmb*+ macrophage to conduct a complete, multi-step lipid-processing cascade: internalization, lysosomal/lipophagic degradation, and efflux. These findings demonstrate that *Gpnmb+* macrophages function as a highly specialized subset regulating ovarian lipid homeostasis.

### Ovarian *Gpnmb+* macrophages interact with lymphatic vessels

Next, we investigated the cellular fate of *Gpnmb+* macrophages following lipid uptake in the regressing CL. Emerging evidence indicates that post-phagocytic macrophages typically undergo apoptosis, migrate via lymphatic vessels, or transition into tissue-resident cells [35, 36]. To evaluate the potential fate of *Gpnmb+* macrophages, we co-immunostained ovaries from mice at 10d post-hCG, targeting GPNMB with either apoptotic marker cleaved Caspase-3 (cl-Caspase-3) or lymphatic vessel marker LYVE1. Whereas robust cl-Caspase-3 signaling was detected within atretic follicles, it did not colocalize with GPNMB (Fig. 7A, left), ruling out apoptosis as the immediate fate of *Gpnmb+* macrophages. In contrast, we observed localization of *Gpnmb+* macrophages adjacent to LYVE1-labeled lymphatic vessels (Fig. 7A, right). To overcome the limitations of 2D tissue sections in capturing cellular interactions, we performed 3D whole-mount staining and imaging on ovaries from mice at d10 post-hCG, which confirmed a direct contact between *Gpnmb+* macrophages and LYVE1+ lymphatic vessels (Fig. 7B).

**Figure 7.**
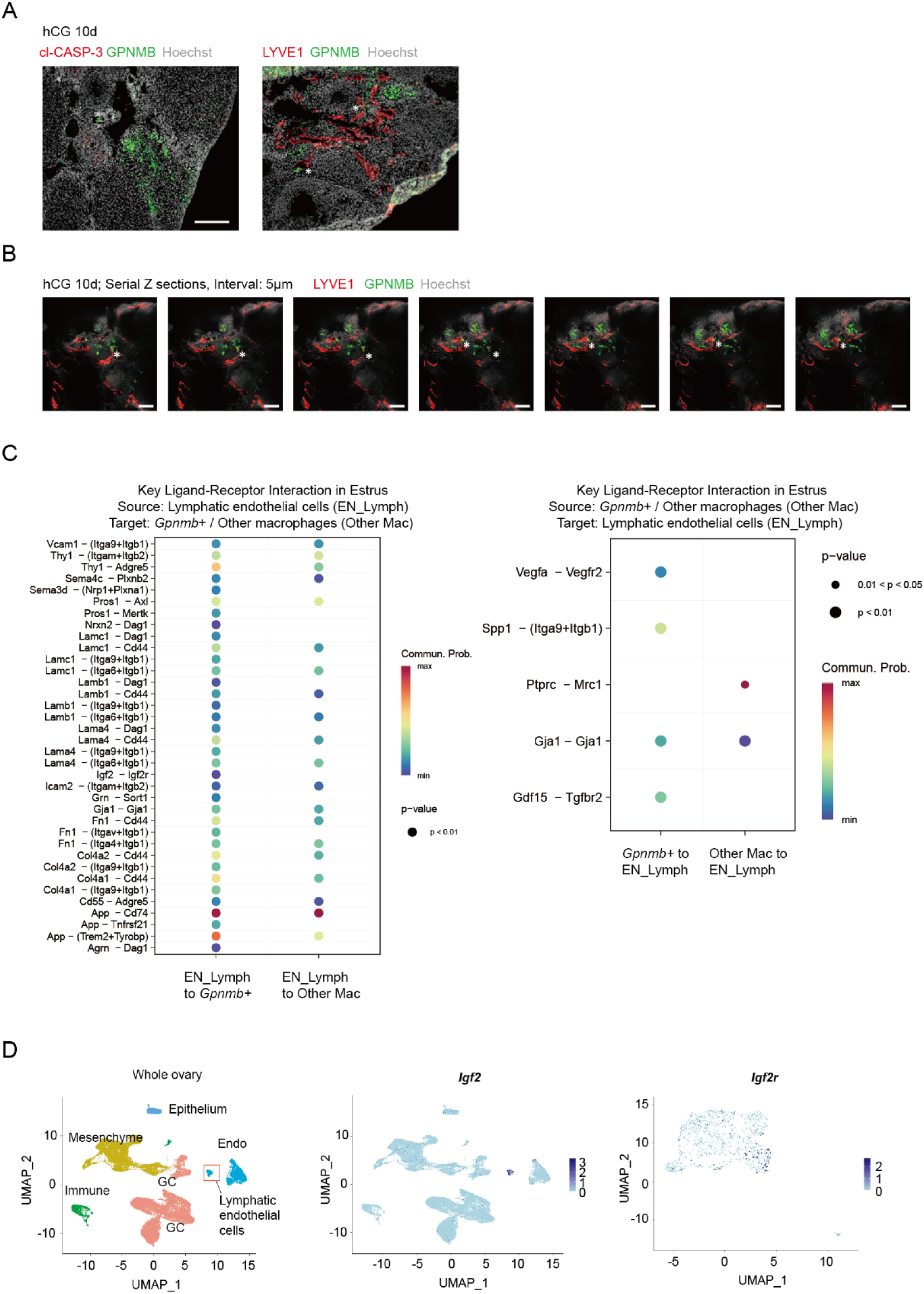
Ovarian *Gpnmb+* macrophages did not undergo apoptosis and were likely to migrate through lymphatic vessel after lipid uptake. (A) Co-immunostaining of GPNMB with cleaved Caspase-3 (cl-CASP-3) (left) or LYVE1 (right) in ovaries collected from mice at d10 post-hCG. Asterisk: *Gpnmb*+ macrophages adjacent to LYVE1+ lymphatic vessels. Scale bar: 200um. (B) Serial Z-sections of whole-mount staining of GPNMB and LYVE1 in ovaries from mice at d10 post-hCG. Asterisk: *Gpnmb*+ macrophages interact with LYVE1+ lymphatic vessels. Scale bar: 100um. (C) Detailed key ligand-receptor interactions between lymphatic endothelial cells and either *Gpnmb*+ macrophages or other ovarian macrophages at estrus, inferred by CellChat analysis [21, 61]. Left: lymphatic endothelial cells to macrophages; right: macrophages to lymphatic endothelial cells. (D) UMAP showing the distribution of *Igf2* transcripts across different ovarian cell types and the distribution of *Igf2r* transcripts across macrophage clusters in the mouse ovary. Data and analyses in C and D were based on an ovarian scRNA-seq dataset [21].

To gain deeper insights into the interaction between lymphatic endothelial cells and ovarian macrophages, we performed cell-cell interaction analysis of ovarian scRNA-seq data [21] focusing on signaling pathways between lymphatic endothelial cells (defined by high transcript levels of *Mmrn1, Ccl21a,* and *Prox1*, Fig. S1A) and *Gpnmb*+ macrophages or other ovarian macrophages. Our analysis revealed several upregulated signaling pathways between lymphatic endothelial cells and *Gpnmb+* macrophages compared to other macrophages. Signaling originating from lymphatic endothelial cells included pathways modulating lipid/phagocytic-sensing programs (*App*-*Trem2/Tyrobp*), cell attachment (*Thy1*-*Adgre5*), cellular clearance (*Pros1*-*Mertk/Axl*), insulin-like growth factor 2 signaling (*Igf2-Igf2r*), and broad extracellular matrix interactions (*Fn1*, *Col4a1/2*, and Laminin axes) (Fig. 7C, left); signaling originating from macrophages indicated that *Gpnmb*+ macrophages mainly drove lymphangiogenesis (*Vegfa*-*Vegfr2*), matrix adhesion (*Spp1*-*Itga9*/*Itgb1*), and stress-associated signaling (*Gdf15*-*Tgfbr2*), while *Gpnmb*-macrophages predominantly engaged in immune-adhesion cross-talk dominated by *Ptprc-Mrc1* (Fig. 7C, right). Notably, *Thy1-Adgre5* signaling pathway is known to facilitate immune cell adhesion to endothelial cells [37, 38], supporting a model where *Gpnmb*+ macrophages exhibit enhanced adhesion to lymphatic endothelial cells. Interestingly, in the mouse ovary, the transcript of *Igf2* is enriched in lymphatic endothelial cells and the transcript of *Igf2r* is enriched in *Gpnmb*+ macrophages compared to other macrophage subsets (Figure 7D). As IGF2 has been reported to reprogram macrophages toward an anti-inflammatory phenotype [39, 40], the cell-type-specific expression of *Igf2* and *Igf2r* in the ovary suggests a role for lymphatic endothelial cells in reducing local inflammation. Taken together, these findings suggest that *Gpnmb+* macrophages may either migrate through lymphatic vessels or directly secrete processed lipids into the lymphatic circulation following lipid engulfment in regressing CL, and lymphatic vessels likely promote the anti-inflammatory phenotype of *Gpnmb*+ macrophages suggested by GO analysis (Fig. 2D).

## Discussion

In this study, we identified a *Gpnmb*+ subset of ovarian macrophages with a distinct transcriptomic profile that is conserved between humans and mice and supports specialized function in phagocytosis, lysosomal processing, and lipid transport. Intriguingly, this gene profile closely mirrors LAMs found in adipose tissue from obese mice and human (Fig. 1D) [18, 41], kidney from obese mice [42], diseased fatty liver [18], and disease-associated microglia in Alzheimer’s Disease in mice [43]. While recent studies have implicated *Gpnmb*+ MNGCs in chronic fibro-inflammatory remodeling in the aging ovary [19] and observed *Gpnmb+* macrophages presence in ovaries of normal cycling mice [20], the function of *Gpnmb*+ macrophages in normal ovarian physiology remained undefined. Our work demonstrates that these *Gpnmb*+ macrophages dynamically expand specifically during luteolysis and physically operate as functional LAMs within a non-pathological context, establishing their essential role in coordinating cyclic lipid homeostasis in the murine ovary.

Immunofluorescence staining demonstrates that in the mouse ovary, *Gpnmb*+ macrophages exclusively reside within regressing CL and surround oocytes in atretic follicles (Fig. 3 & 4). As luteal cells and oocytes contain higher lipid content compared to other cell types in the murine ovary [12–14, 44], this restricted localization of *Gpnmb*+ macrophages aligns with previous findings of LAMs in adipose tissue, where they accumulate markedly more lipid droplets than other resident macrophage subsets [41]. As we observed only few *Gpnmb*+ macrophages in the ovary outside atretic follicles before d3 post-hCG, it is plausible that after precursor monocytes/macrophages are recruited into the regressing CL, they undergo localized differentiation in response to high lipid content or other molecular cues. However, the exact mechanism driving this process remains to be fully elucidated. Regressing CL are known to secrete signals to recruit immune cells [45]. In agreement with this, our ligand-receptor interactome analysis revealed possible recruiting signals between regressing luteal cells and *Gpnmb*+ macrophages, including Semaphorin pathways [46] and the *Cxcl12*-*Cxcr4* axis (Fig. 5). Future studies are warranted to explore the importance of these pathways in recruiting monocytes/macrophages during structural luteolysis.

Given that ovarian *Gpnmb*+ macrophages shared a similar gene profile with LAMs, which fail to differentiate upon the knockout of *Trem2* [18], it is highly probable that the differentiation of ovarian *Gpnmb*+ macrophages is also *Trem2*-dependent. Consequently, evaluating ovarian lipid clearance capacity using a *Trem2*-KO mouse model represents an exciting future direction. TREM2 is an innate immune receptor that acts as a broad-spectrum sensor for tissue damage, cellular debris, and pathogens [47]. Although the abundant cellular debris within regressing CL may readily activate TREM2 to initiate cell differentiation, this alone may be insufficient for the differentiation of *Gpnmb*+ macrophages, as we only observed few *Gpnmb*+ macrophages in atretic follicles in immature mice and they are mostly associated with degenerating oocytes (Fig. 4D). Given that lipid processing is a core function of *Gpnmb*+ macrophages, localized lipids are likely to serve as a necessary co-activator for their differentiation. Regressing CL are full of lipid droplets containing triglyceride and cholesterol [12–14]; and oocytes, the largest cells in the body, have a high density of internal lipid droplets [44]. Therefore, we propose a model wherein the sensing of localized lipid loads and apoptotic debris released from dying cells are required to fully activate the differentiation of *Gpnmb*+ macrophages.

While a previous study in non-human primates indicated that luteal cells themselves can efflux cholesterol via ABCA1 during luteolysis [15], whether immune cells actively execute reverse cholesterol transport as a dedicated clearance mechanism in regressing CL remained to be determined. Transcriptomic profile of *Gpnmb*+ macrophages is enriched in genes related to reverse cholesterol efflux (*Abca1* and *Abcg1*), closely mirroring the upregulated *Abca1/Acbg1* in foam cells in the arterial walls of atherosclerosis patients [48], and LAMs in kidney of obese mice [42]. In foam cells, intracellular cholesteryl esters are hydrolyzed and exported as free cholesterol via ATP-binding cassette transporters (ABCA1 and ABCG1), and scavenger receptor class B member 1 (SR-BI) [49, 50]. It is plausible that *Gpnmb*+ macrophages utilize similar strategies to clear the excess ovarian cholesterol. Based on the role of lymphatic vessels in transporting lipids following reverse cholesterol transport [51] and during luteolysis [52], our findings demonstrating direct physical contacts between *Gpnmb*+ macrophages and lymphatic endothelial cells provide strong support for a macrophage-lymphatic vessel-mediated lipid resolution pathway during luteolysis. Ultimately, this coordinated axis of macrophage-mediated reverse cholesterol transport and lymphatic drainage, similar to the cholesterol efflux mechanisms observed in foam cells [48], highlights a highly specialized, non-inflammatory mechanism for lipid homeostasis in the ovary following luteal regression.

Following phagocytosis, macrophages typically undergo apoptosis, migrate via lymphatic vessels, or transition into tissue-resident cells [35, 36]. In our analysis, we ruled out immediate local apoptosis, as we did not detect apoptotic markers within the *Gpnmb*+ cells by day 10 post-hCG (Fig. 7A). In addition to direct contact between lymphatic vessels and *Gpnmb+* macrophages (Fig. 7B), the enhancement of cell adhesion pathway (*Thy1-Adgre5 and Spp1-Itga9/Itgb1*) between *Gpnmb*+ macrophages and lymphatic endothelial cells supports their active and direct interaction (Fig. 7C). While our data support the hypothesis that *Gpnmb+* macrophages migrate through lymphatic vessels, we cannot exclude the possibility that some of them may differentiate into ovarian resident macrophages. Definitive evidence to distinguish these possibilities will require future high-resolution cell-tracing studies. Additionally, determining the ultimate cell fate and recruitment dynamics of LAMs remains a fundamental challenge in metabolic research [53]. Therefore, beyond advancing our understanding of reproductive biology, we suggest that further studies in mouse ovaries can provide informative insights into possibly conserved mechanisms related to metabolic homeostasis.

Elucidating the function of *Gpnmb*+ macrophages may also provide insights into understanding ovarian aging. MNGCs have been found in aged ovaries in mice and non-human primates [54, 55]. Existing evidence suggest that MNGCs develop through the fusion of macrophages and are often present in chronic inflammatory conditions, including infectious or non-infectious granulomas [56]. Intriguingly, a recent study indicates *Gpnmb* is a definitive marker of these giant cells [19]; therefore, it is highly plausible that MNGCs in aged ovaries are derived from *Gpnmb*+ macrophages from previous cycles. Current theories suggest that MNGC formation in the aging ovary is driven by an overwhelming accumulation of metabolic waste and cellular debris that individual macrophages cannot efficiently degrade, forcing them to fuse [19]. Moreover, recent evidence indicates that aging ovarian macrophages also undergo an intrinsic decline in phagocytic and degradative capacity [20], likely propelling them to fuse into MNGCs as a compensatory response. Future studies are warranted to delineate the lineage and functional relationship between *Gpnmb*+ macrophages and MNGCs.

Impaired lipid clearance and subsequent accumulation within the ovary can lead to lipotoxicity, a pathological state related to various female reproductive diseases [57, 58]. For instance, high-fat diet-induced lipid accumulation in the ovary can lead to reduced mitochondrial membrane potential in the oocyte and endoplasmic reticulum stress in cumulus cells [59]. Additionally, an elevated abundance of lipid droplets within granulosa cells is associated with a decline in the rate of successful conception among IVF patients [60]. Collectively, these physiological and clinical observations underscore the importance of maintaining lipid homeostasis in the ovarian microenvironment. Functional impairment or failure of *Gpnmb*+ macrophages to clear lipids from regressing CL may accelerate ovarian aging and compromise fertility via chronic lipotoxicity. While our current study provides strong observational and transcriptomic evidence for the lipid uptake capability of ovarian *Gpnmb*+ macrophages, a key limitation is the lack of direct loss-of-function experiments. Future studies utilizing tissue-or cell-type-specific genetic knockout mouse models or targeted depletion strategies will be essential to directly test the requirement of *Gpnmb*+ macrophages in maintaining cyclic ovarian homeostasis, as well as how their dysregulation may contribute to pathological changes in the ovary.

In conclusion, we have characterized a distinct *Gpnmb*+ macrophage subset in the normal cycling ovary that exhibits a gene expression signature closely resembling LAMs in other pathological conditions. Functionally, these macrophages are responsible for taking up and clearing lipids during CL regression. Collectively, these data expand the knowledge of LAM biology and ovarian physiology, demonstrating that *Gpnmb*+ macrophages are not restricted to pathological states of chronic inflammation and age-related degeneration but are finely tuned mediators of normal ovarian homeostasis.

**Figure S1.**
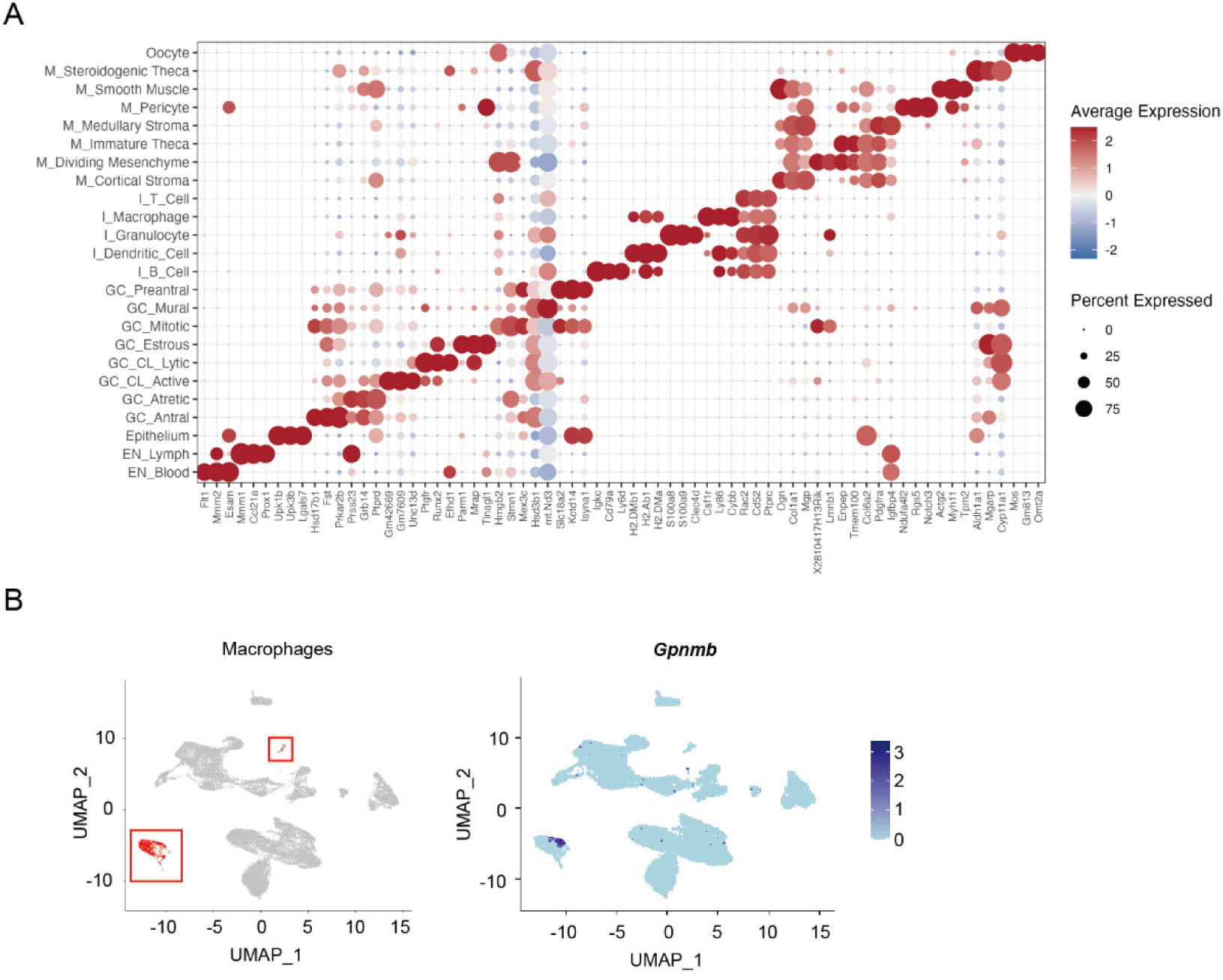
Marker gene expression for the cell type annotations of the cycling mouse ovary dataset used for downstream analyses. (A) Dot plot showing the three most enriched marker genes for each annotated cell population in the cycling mouse ovary scRNA-seq dataset from Morris *et al.*, 2022 [21]. GC, granulosa cell; M, mesenchyme; I, immune; EN, endothelium; CL, corpus luteum. (B) UMAP showing that the expression of *Gpnmb* is restricted in macrophages in the mouse ovary.

**Figure S2.**
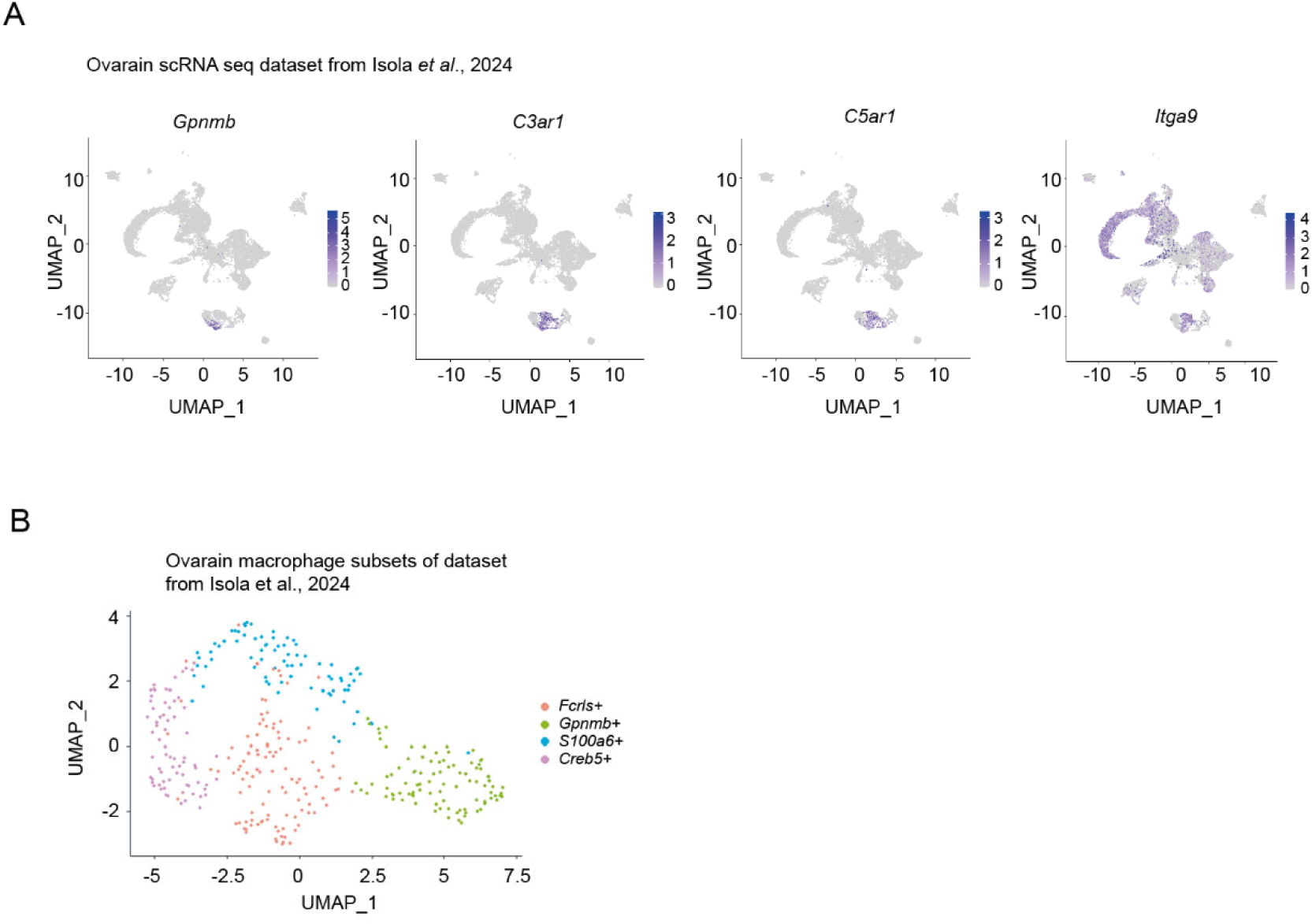
The *Gpnmb+* macrophages subpopulation is confirmed in an additional mouse ovarian scRNA-seq dataset. (A) UMAP showing the pattern of expression of *Gpnmb, C3ar1, C5ar1,* and *Itga9* cross different cell clusters within an another mouse ovarian scRNA-seq dataset (both ages, 3 months and 9 months) [22]. (B) UMAP of macrophage subsets in the same ovarian scRNA-seq dataset.

**Figure S3.**
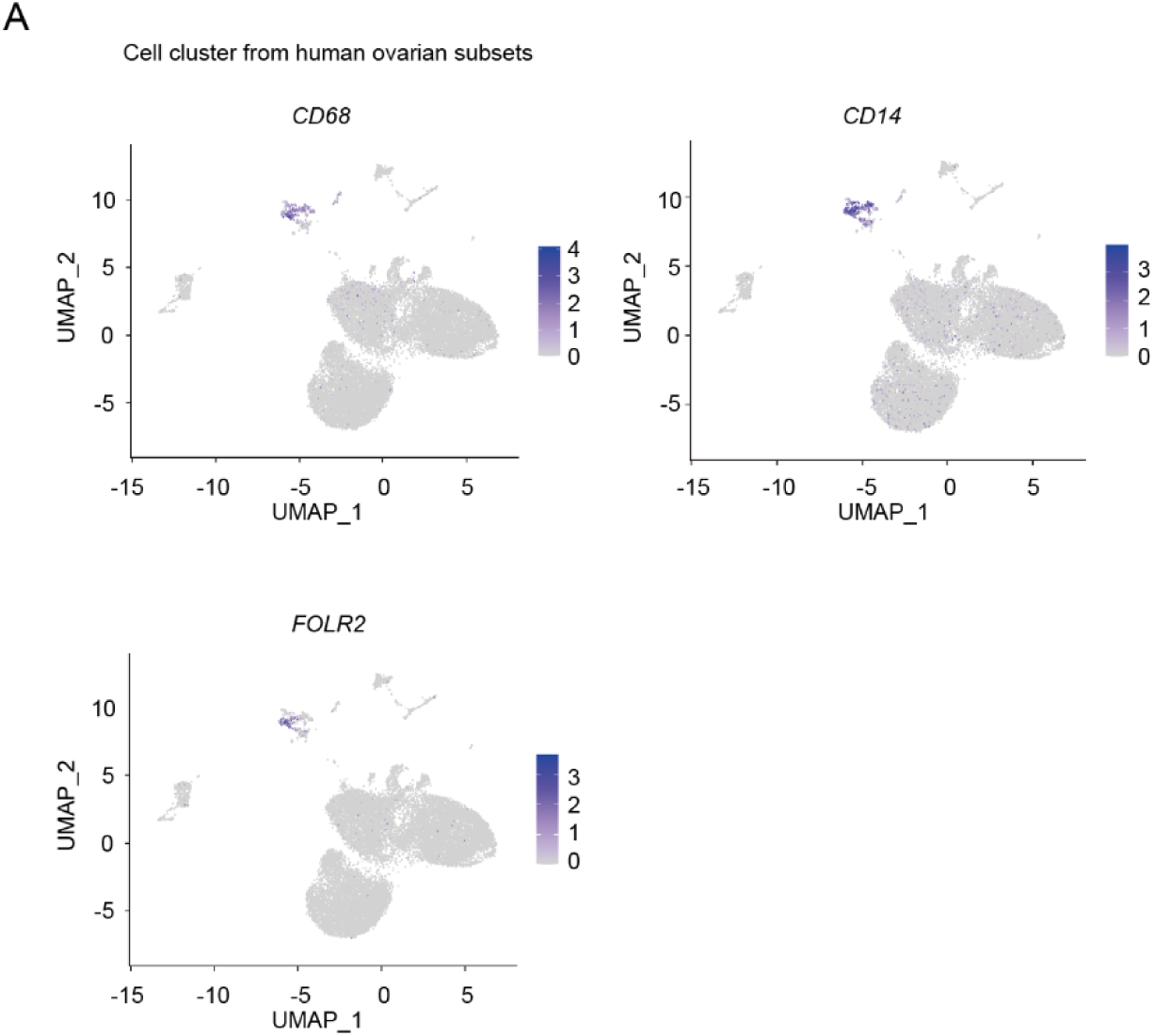
The marker genes of macrophages in a human ovarian scRNA-seq dataset. (A) UMAP showing the expression of *CD68, CD14,* and *FOLR2* across different cell clusters in a human ovarian scRNA-seq dataset [23].

**Figure S4.**
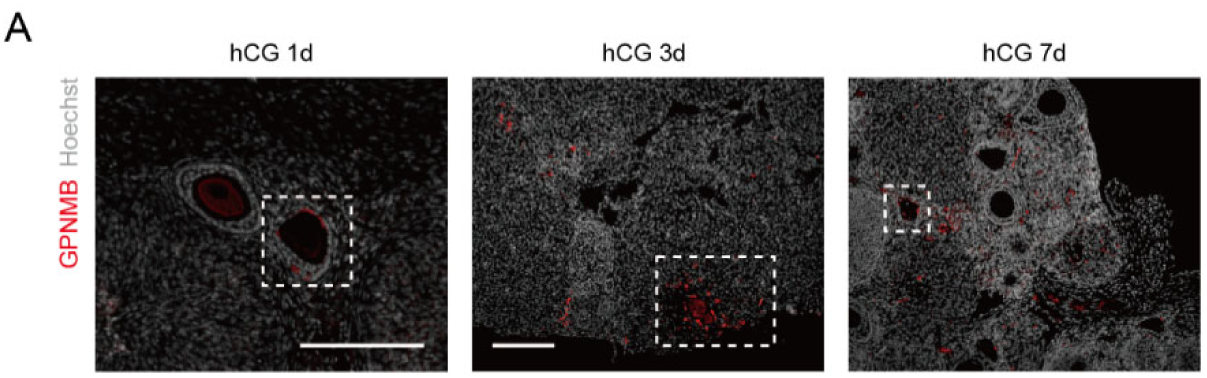
*Gpnmb*+ macrophages surround oocytes at different time points post-hCG. (A) Immunostaining of GPNMB in ovaries collected from d1, 3, and 7 post-hCG showing that *Gpnmb*+ macrophages surround oocytes. Scale bar: 200um.

**Figure S5.**
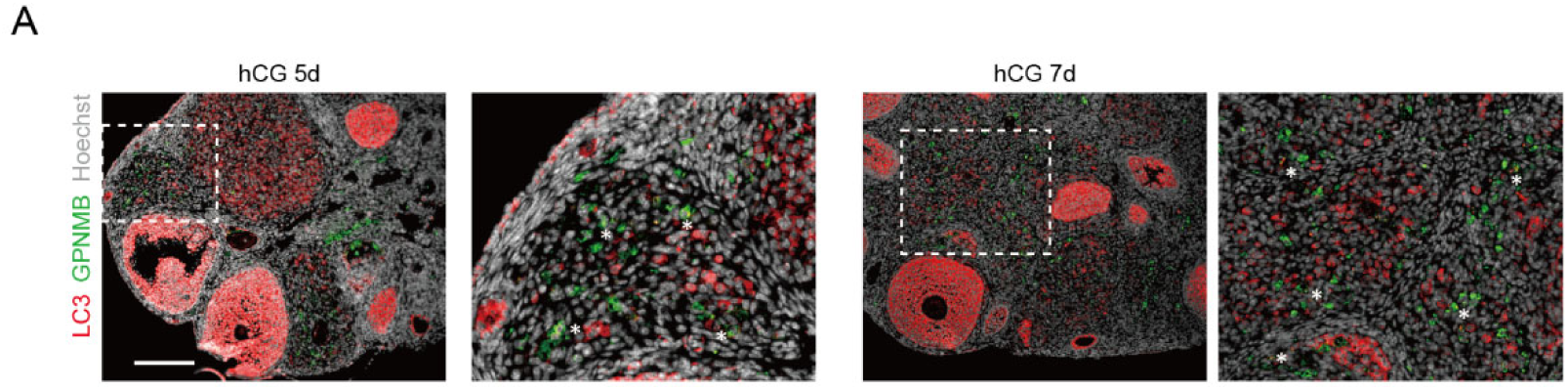
Ovarian *Gpnmb+* macrophages likely utilize lipophagy for lipid clearance. (A) Immunostaining of GPNMB and LC3 in ovaries collected from mice at d5 and 7 post-hCG. Asterisk: *Gpnmb*+ macrophages with LC3 puncta. Scale bar: 200um.

## Materials and Methods

### Animals

Experiments were performed using C57BL/6J mice generated from our in-house breeding colony. Animals were maintained in individually ventilated polycarbonate cages (11.5 * 7.5 inches) under controlled environmental conditions (68-77℉, 30-70% relative humidity) with a standard 14-hour light/ 10-hour dark cycle. All experimental protocols strictly adhered to the U.S. Public Health Service Policy on Humane Care and Use of Laboratory Animals and were formally approved by the Institutional Animal Care and Use Committee (IACUC) at Cornell University under protocol #2019-0006. To minimize distress during euthanasia, mice were humanely euthanized via CO_2_ inhalation, administered at a chamber displacement rate of 30-70% per minute until full cessation of respiration and deep unconsciousness were confirmed. Cervical dislocation was subsequently executed as a secondary physical method to guarantee death.

### Hormone stimulation of superovulation model

To induce superovulation, prepubertal female mice (aged 21-23 days, body weight 10-11 g) were administered a single intraperitoneal (IP) injection of 5 IU pregnant mare serum gonadotropin (PMSG; NATE-0969, Creative Enzymes). Ovulation was subsequently triggered 44-48 hours post-PMSG by an IP injection of 5 IU human chorionic gonadotropin (hCG; 9002-61-3, Sigma-Aldrich).

### RNA extraction, reverse transcription, and real-time quantitative PCR (RT-qPCR)

Total RNA was isolated using the RNeasy Micro Kit (74106, Qiagen) according to the manufacturer’s instructions. Complementary DNA (cDNA) was synthesized using the High-Capacity cDNA Reverse Transcription Kit (4368814, Applied Biosystems). Quantitative real-time PCR (RT-qPCR) assays were conducted with RT^2^ SYBR Green ROX FAST Mastermix (330623, Qiagen) on a StepOnePlus Real-Time PCR System (4376600, Applied Biosystems). Target gene expression levels were quantified using the 2^(-ΔCt)^ method, with the ribosomal gene *Rpl19* serving as the internal reference control. Specific primer sequences used in this study were as follows: *Gpnmb* (Forward: AGAAATGGAGCTTTGTCTACGTC, Reverse: CTTCGAGATGGGAATGTATGCC); *Trem2* (Forward: CTGGAACCGTCACCATCACTC, Reverse: CGAAACTCGATGACTCCTCGG); *Atp6v0d2* (Forward: CAGAGCTGTACTTCAATGTGGAC, Reverse: AGGTCTCACACTGCACTAGGT); and *Rpl19*(Forward: GGTGACCTGGATGAGAAGGA, Reverse: TTCAGCTTGTGGATGTGCTC).

### Immunofluorescence (IF) staining

Freshly dissected ovaries were immediately embedded in Optimal Cutting Temperature (OCT) compound, and cryosectioned into 7-μm-thick slices for histological analysis or immunofluorescence (IF) staining. Sections designated for IF were fixed in pre-cooled methanol for 10 min at -20°C. Nonspecific binding was blocked by incubating sections in phosphate-buffered saline (PBS) containing 0.1% Tween 20 and 2% normal donkey serum (NDS) for 1 h at room temperature. Tissues were subsequently incubated overnight with primary antibody at 4°C (anti-GPNMB, AF2330, R&D systems, 1:200; anti-cl-Caspase-3, 9661S, Cell Signaling Technology, 1:200; anti-F4/80, 14-4801-82, Invitrogen, 1:50; anti-LAMP1, 14-1071-81, Invitrogen, 1:200; anti-LC3, 12741S, Cell Signaling Technology, 1:200; anti-LYVE1, ab14917, Abcam, 1:200). Following PBS washes, sections were incubated with secondary antibody (Donkey Anti-Rabbit IgG (H+L) conjugated with Alexa 594, A-21207, Invitrogen; Donkey Anti-Goat IgG (H+L) conjugated with Alexa 647, A-32849, Invitrogen; Donkey Anti-Rat IgG (H+L) conjugated with Alexa 594, A-21209, Invitrogen) at 1:1000 dilution for 1 h at room temperature. Nuclei were counterstained with 1 μg/ml Hoechst 33342 (76482-876, VWR). For neutral lipid labeling, rinsed sections were probed with BODIPY (1μM) for 20 min at room temperature. Fluorescence images were captured using a Nikon Diaphot 300 microscope (Nikon Instruments) and post-processed using ZEISS ZEN and ImageJ software.

### Whole-mount staining

Ovaries harvested from immature and adult mice were fixed in 4% paraformaldehyde for 4 h and subsequently stored in 70% ethanol. Fixed ovarian tissues were cut into 3-4 pieces and progressively rehydrated in PBS. Heat-induced antigen retrieval was carried out at 70°C for 30 min in 10 mM sodium citrate + 0.05% v/v Tween-20, pH 6.0. Tissues were permeabilized with PBS containing 1% Triton X-100 for 30 min at room temperature and blocked in PBS containing 0.1% Triton X-100 and 5% normal donkey serum (NDS) for 1 h at room temperature. Samples were incubated with antibodies in PBS containing 0.1% Triton X-100 and 2% NDS (PBS-TX-NDS) overnight at 4°C (anti-GPNMB, AF2330, R&D systems, 1:50; anti-LYVE1, ab14917, Abcam, 1:50). After rinsing, samples were incubated with secondary antibody (Donkey Anti-Rabbit IgG (H+L) conjugated with Alexa 594, A-21207, Invitrogen; Donkey Anti-Goat IgG (H+L) conjugated with Alexa 647, A-32849, Invitrogen) at 1:100 dilution overnight at 4°C plus 2 μg/ml Hoechst 33342 (76482-876, VWR) in PBS-TX-NGS. For the labeling of neutral lipids, rinsed samples were probed with BODIPY (2μM) for 30 min at room temperature. After rinsing, tissues were mounted in Aqua-Poly/Mount (Polysciences Inc.) in coverwell imaging chambers (70327-08, Electron Microscopy Sciences). Ovaries were imaged using an inverted laser scanning confocal microscope (Zeiss LSM880 microscope). Images were taken at intervals (3 or 5µm) in z-stack starting from the outer surface of the CL and processed with ZEISS ZEN.

### Bioinformatics and public dataset analysis

Publicly available scRNA-seq datasets of the cycling mouse ovary (Morris *et al.*, 2022 [21], Broad Institute Single Cell Portal ID SCP1914), aging mouse ovary (Isola *et al*., 2024 [22]; GEO: GSE232309), and healthy human ovary (Jones *et al*., 2024 [23]; GEO: GSE260686), were analyzed using the Seurat R package (v 5.3.0). For the cycling mouse ovarian dataset, we used the cell type annotations in the metadata from the original study [21]. To confirm these annotations, marker genes were identified using the FindAllMarkers function with a Wilcoxon rank-sum test, retaining positively enriched genes (min.pct = 0.25, |log_2FC| > 0.25, adjusted *p*<0.05). The identified markers were provided in Figure S1. For other datasets, ovarian macrophages were annotated based on canonical markers used by each study (*C3ar1*/*Itga9*/*C5ar1* for aging mouse [22]; *CD68*/*CD14*/*FOLR2* for human [23]). For reclustering, briefly, gene expression data were normalized using the LogNormalize method, and top variable features (typically top 2,000) were identified for data scaling and linear dimensionality reduction via Principal Component Analysis (PCA). To capture population heterogeneity across specific cell lineages and conditions, graph-based clustering (FindNeighbors and FindClusters) and Uniform Manifold Approximation and Projection (UMAP) were performed using dataset-optimized principal components (e.g., top 7-30 PCs) and resolution parameters (typically 0.2-0.8). Differentially expressed genes (DEGs; adjusted *p*<0.05, |log_2FC| > 0.25) were identified via Seurat, and functional pathway enrichments for Gene Ontology Biological Process (GO-BP) and KEGG pathways were determined using clusterProfiler (v4.20.0). Human gene symbols were converted to lowercase to align with mice ortholog nomenclature prior to the intersect analysis [62]. The overlapping markers between human and mice was evaluated using hypergeometric testing (phyper) [62].

To infer stage-specific cell-cell communications during estrus, ligand-receptor interaction analyses between *Gpnmb*+ macrophages and active or dying luteal cells were performed using CellChat (v2.2.0.9001), following the standardized protocol described by Jin *et al*. [61]. To examine the spatial distribution of *Gpnmb* in the ovary, a publicly available spatial transcriptomic dataset of the ovulating mouse ovary from Mantri *et al*., 2024 was used [24] (GEO: GSE240271). The processed anndata file was downloaded from figshare, and normalized *Gpnmb* expression was visualized for each timepoint using scanpy (v1.9.1) and squidpy (v1.2.3). Cell type annotations were the same as in the original publication. Furthermore, to assess transcriptional similarity between *Gpnmb*+ macrophages from the Morris *et al*., 2022 paper and LAMs described by Jaitin *et al*., 2019 [18] (GEO: GSE128518), differential expression profiles between the two datasets were compared. First, genes differentially expressed in *Gpnmb*+ macrophages relative to other macrophages from the ovary were identified using the wilcoxon rank-sum test in Seurat (v5.3.0) and the log2 fold changes of these DEGs were calculated. Next, log2 fold changes were obtained from Jaitin *et al.*, 2019, corresponding to the comparison of Mac3 (LAM) macrophages from high-fat-diet mice against Mac1 (homeostatic) macrophages from normal-weight mice. Genes present in both datasets and the corresponding log2 fold changes were compared by a Pearson correlation test. Correlations were visualized as scatter plots, with canonical LAM markers labelled using ggplot2 (v4.0.2) and ggrepel (v0.9.6).

## Statistical analysis

All quantitative data are presented as mean ± standard deviation (SD). Statistical analysis was performed using the GraphPad Prism 10 analysis software. Each experiment included at least three independent biological samples and was repeated at least three times. For comparison across three or more groups, One-way ANOVA followed by Tukey’s multiple comparisons test was used. *p* < 0.05 was considered statistically significant.

## Declaration

## Acknowledgments

This work was supported by R01HD109392 from the NICHD (to Y.A.R.).

## Consent to participate

Not applicable.

## Consent for publication

Not applicable.

## Ethical approval and its number

All animal procedures were conducted in accordance with the U.S. Public Health Service Policy on Humane Care and Use of Laboratory Animals and were approved by the Institutional Animal Care and Use Committee at Cornell University (IACUC protocol #2019-0006).

## Data availability

All datasets analyzed in this study are publicly available in open repositories: cycling mouse ovarian scRNA-seq data from Broad Institute Single Cell Portal (SCP1914) and OSF (924fz); aging mouse ovarian scRNA-seq (GSE232309), healthy human ovarian scRNA-seq (GSE260686), preovulatory spatial transcriptomics (GSE240271), and adipose LAM scRNA-seq (GSE128518) from NCBI Gene Expression Omnibus (GEO).

## Author contribution

Y.H.L. and Y.A.R. conceived of and designed the study. Y.H.L. conducted experiments. Y.H.L. and K.J. analyzed the data. All authors contributed to the manuscript and provided feedback and comments.

